# Full-length isoform constructor (FLIC) – a tool for isoform discovery based on long reads

**DOI:** 10.1101/2025.05.27.656444

**Authors:** Alexandra M. Kasianova, Anna V. Klepikova, Oleg A. Gusev, Guzel R. Gazizova, Maria D. Logacheva, Aleksey A Penin

## Abstract

Advances in high-throughput sequencing have illuminated a complexity of transcriptome landscape in eukaryotes. An inherent part of this complexity is the presence of multiple isoforms generated by the alternative splicing and the use of alternative transcription start and polyadenylation sites. However, currently available tools have limited capacity to infer full-length isoforms. To address this problem, we developed a new pipeline, FLIC (Full-Length Isoform Constructor). FLIC is based on the long-read transcriptome data and integrates several key features: 1) utilizing biological replicate concordance to filter out noise and artifacts; 2) employing peak calling to precisely identify transcription start and polyadenylation sites; 3) enabling robust isoform reconstruction with minimal reliance on existing annotations. We evaluated FLIC using a dedicated set of real and simulated data of Arabidopsis thaliana cDNA sequencing. Results demonstrate that FLIC accurately reconstructs known and novel isoforms, outperforming existing tools, especially in the absence of reference annotations. A direct comparison with CAGE, currently regarded as golden standard for TSS identification shows that FLIC is equally accurate, while being much less time-consuming. Thus FLIC provides a valuable tool for comprehensive transcript characterization, particularly for non-model organisms or when dealing with incomplete or inaccurate annotations.

## 1 Introduction

Splicing is a key molecular event in mRNA processing. Its regulation, including the generation of alternative isoforms plays crucial role in cell differentiation, developmental changes, environmental adaptations (stress response) and pathogen response in animals and plants (e.g. (Dikaya *et al*., 2021; Rotival *et al*., 2019). Splicing has been shown to vary not only between tissues, organs and developmental stages but also between individuals and populations (Park *et al*., 2018). The deeper understanding of splicing is important for biomedical applications because alterations of splicing events are hallmarks (and in some cases, drivers) of many pathological conditions such as cancer and neurodegenerative diseases (reviewed in (Nikom and Zheng, 2023; Bradley and Anczuków, 2023).

The advent of high-throughput sequencing equipped the researchers with new tools for the study of splicing. Second-generation platforms such as Illumina and MGI, allow simultaneous characterization of splice sites in all genes expressed in a certain sample. Being able to provide accurate genome-wide quantification of splicing sites, short-read technologies however have limited capacities in characterization of full-length isoforms. This challenge is addressed in 3rd generation sequencing platforms, namely SMRT from Pacific Biosciences and nanopore sequencing (Oxford Nanopore Technologies) which allow sequencing full-length transcripts (for review see (De Paoli-Iseppi *et al*., 2021). However they have their drawbacks (high error rate, incomplete reading of transcripts); several tools were developed to mitigate it but the problem is far from being solved.

We propose a new approach based on the long transcriptome reads that is centered on experimental data and is able to accurately identify known and novel transcripts.

## 2 Materials and Methods

### 2.1.1 Plant growing and RNA extraction

*Arabidopsis thaliana* plants of Col-0 ecotype were grown in a climate chamber (16 hours light/ 8 hours dark, 22C, 60% relative humidity) for 21 days after germination. Mature leaves (pools taken from five plants) were collected in three replicates. For RNA extraction we used RNAeasy Plant Mini kit (Qiagen, Venlo, The Netherlands) according to manufacturer’ instructions. RNA quality was accessed using Bionalyzer 2100 (Agilent, Santa Clara, CA, USA) run in Plant RNA mode. RIN was higher than 8 for all samples.

### 2.1.2 ONT and CAGE library preparation

For ONT libraries RNA was converted to cDNA using Mint cDNA synthesis kit (Evrogen, Moscow, Russia) with 18 cycles of amplification with the following modification: 1) cDNA synthesis is primed with oligonucleotides that contain not only dT part and adapter part but also custom barcode, specific for each sample; 2) optimal number of cycles for amplification was selected based on the results of qPCR with primers annealing at 3’ and 5’ ends. The number of cycles falling to the upper part of linear phase was considered to be optimal. Amplified cDNA was used as input for library preparation using standard protocol for genomic DNA with LSK-109 kit. Libraries were sequenced on PromethION instrument using R9.4.1 flowcell. For CAGE, we used the same RNA and processed it as described in Murata et al. (Murata *et al*., 2014).

### 2.2.1 ONT read processing

Basecalling was performed by Guppy 3.4.5 (Oxford Nanopore Technologies, Oxford, United Kingdom). After that, reads were processed by a custom pipeline NTproc (https://github.com/shelkmike/NTproc/, (Kasianova *et al*., 2024). It removes the reads that do not have adapter sequences on both ends and converts reads that had barcodes on their 5’-ends in order to unify the orientation of the reads (Supplementary Fig. 1B).

### 2.2.2 Trimming of M1-filtered ONT reads

PolyA tails were removed from M1-filtered ONT reads using cutadapt v. 4.4 (Martin, 2011) with parameter “*-a A{100}*”.

### 2.2.3 Trimming of Illumina reads for the splice site identification

Raw RNA-seq Illumina reads were trimmed using Trimmomatic v. 0.36 (Bolger *et al*., 2014) with following parameters: “*SE -phred33 ILLUMINACLIP:adapters_filename:2:30:10 LEADING:20 TRAILING:20 SLIDINGWINDOW:4:15 MINLEN:30*”.

### 2.2.4 Trimming of CAGE data

Illumina reads obtained from the CAGE libraries were trimmed using fastp v. 0.23.4 (Chen *et al*., 2018) with parameters “*--cut_right --cut_front --trim_tail1 1*”.

### 2.2.5 Mapping of CAGE data

Trimmed CAGE reads were mapped to the reference genome assembly (*A. thaliana* TAIR10.1) using bwa-mem2 v. 2.2.1 (Vasimuddin *et al*., 2019) with default parameters.

### 2.2.6 Mapping of ONT reads

ONT reads were aligned to the reference genome assembly (*A. thaliana* TAIR10.1) using minimap2 v. 2.24 (Li, 2018) with the following parameters: *-ax splice -k14 -uf -G 10k*. A custom script was used to select reads with a single alignment only.

### 2.2.7 Generation of Illumina splice site reference dataset

A splicing site database was generated by mapping of trimmed RNA-seq Illumina reads to a reference genome assembly (*A. thaliana* TAIR10.1) with STAR v. 2.7.10b (Dobin *et al*., 2013) using the following parameters: “*--outFilterMismatchNmax 2 --outSJfilterCountUniqueMin 3 1 1 1 1 --outSJfilterCountTotalMin 3 1 1 1 1 --alignIntronMin 15*”. Only splice junctions that appeared in two or more samples were considered.

### 2.2.8 Splice sites identification and correction

For each read, the coordinates of extended gaps labeled with the symbol “N” were extracted from the CIGAR string of an alignment and stored in an intermediate file. If coordinates of a given gap in a read were within 10 bp range of any known splice site from the Illumina splice site reference dataset, the coordinates of a gap are replaced by the coordinates of a corresponding splice site from a reference dataset in the intermediate file. Otherwise, an extended gap is considered incorrectly identified and excluded from the description of the read.

### 2.2.9 Read downsampling of highly expressed genes

Reads downsampling based on TAIR10.1 genome annotation was performed using the optional custom module with default parameters. Only genes covered with at least five ONT reads were considered. Genes with more than 1 000 (by default) ONT reads mapped were subjected to downsampling: for each gene 1 000 ONT reads were randomly selected from all reads mapped to a given gene.

### 2.2.10 Processing of mapping data for peak calling

For ONT data read alignments in SAM format were converted to BAM format using the samtools view v. 1.15.1 (Danecek *et al*., 2021) with the parameters “*-Sb*”.

For CAGE data read alignments in SAM format were converted to BAM and only primary mapped reads were retained using samtools view v. 1.15.1 (Danecek *et al*., 2021) with the following parameters: “*-Sb -F 0×100 -F 0×800 -F 0×4*”.

Next, for both data types bam files were sorted using samtools sort. Bedtools v. 2.30.0 program (Quinlan and Hall, 2010) were used to estimate the coverage by 5’- or 3’-ends of ONT reads separately for the forward and reverse strands with the following command: *bedtools genomecov -[5*|*3] -strand [+*|*-] --bg*. The resulting bedGraph files were converted with a custom script in a single-nucleotide format of covered positions. Next, BedGraphToBigWig tool v. 4 (https://github.com/ENCODE-DCC/kentUtils/) was used to convert bedGraph to bigWig format.

### 2.2.11 Peak calling for the detection of TSSs and PAs

BigWig files were processed by CAGEfightR R package (Thodberg *et al*., 2019). No TPM normalization was applied and the following parameters were used: positions with zero coverage were considered “unexpressed” in a *calcSupport* function, then, only positions that were covered in two or more replicates (*support > 1* in a *subset* function). Peak calling was performed with a function “*clusterUnidirectionally*” and parameters *pooledCutoff = 2, mergeDist = 10*.

### 2.2.12 Isoforms and genes reconstruction

Custom script was used to associate each read with TSS and PA regions identified by peak calling. Reads which 5’- or 3’-ends did not fall into any TSS or PA regions were excluded from the analysis. Isoforms were formed by reads with identical TSS, splice sites and PA, and the number of reads forming each isoform was counted. Only isoforms with at least five reads in one replicate and at least one read in another replicate were retained. To form genes out of isoforms the intersection of all vs all isoforms was calculated: the length of an isoform was determined as “right coordinate minus left coordinate”, and if 80% of length of the shorter isoform was intersected with another isoform, these isoforms were combined into a gene. Gene coordinates were determined as most left and most right coordinates of intersected isoforms. Gene IDs were assigned to genes and isoforms according to the *Arabidopsis* genome annotation if reconstructed gene was intersected with a gene from annotation in at least 80% of length of the shorter gene. Next, isoforms belonged to a gene were filtered as followed: the isoform formed by the highest number of ONT reads was declared major, and other isoforms were retained if the number of their reads was higher than 1% of major isoform reads.

### 2.2.13 Selection of genes expressed in mature *Arabidopsis* leaf

To estimate gene expression level in mature *Arabidopsis* leaves, NCBI SRA (https://www.ncbi.nlm.nih.gov/sra) accessions SRR3581681 and SRR3581847 were used. Raw RNA-seq Illumina reads were trimmed using Trimmomatic v. 0.36 (Bolger *et al*., 2014) with following parameters: “*SE -phred33 ILLUMINACLIP:adapters_file_name:2:30:10 LEADING:20 TRAILING:20 SLIDINGWINDOW:4:15 MINLEN:30*”. Trimmed reads were aligned to the reference *A. thaliana* (TAIR 10.1) genome and read number per genes were counted using STAR with the following parameter: *“--quantMode GeneCounts*”. Gene expression was normalized between samples on library size with DESeq median method (Love *et al*., 2014). Genes which expression were higher than 16 normalized readcounts in each sample were selected as expressed.

### 2.2.14 Read simulation

Reads were simulated in three replicates using Arabidopsis thaliana TAIR10.1 reference genome and IsoSeqSim v. 0.2 software (https://github.com/yunhaowang/IsoSeqSim) with the following parameters: “-nbn 1 --nbp 0.004 --c5 5_end_completeness.PacBio-P6-C4.tab --c3 3_end_completeness.PacBio-P6-C4.tab”.

### 2.2.15 Generation of distorted annotation

In order to produce distorted annotation modeling the annotation which contains errors in prediction on 5’- and 3’-ends and intron/exon boundaries, we modified TAIR10.1 annotation in the following way. First, we retained in the GFT file only transcripts coding mRNAs (52 059 transcripts in total). Then using a custom script we randomly changed the boundaries of transcripts in one of the ways: 5’-end moved 100 bp downstream, 5’-end moved 100 bp upstream, 3’-end moved 100 bp downstream, 3’-end moved 100 bp upstream, 5’- and 3’-ends moved 100 bp upstream, 5’- and 3’-ends moved 100 bp downstream or no changes.

### 2.2.16 Transcript reconstruction based on simulated reads

For the experiment on simulated reads isoforms were reconstructed with FLIC v. 1.0 and three other state-of-the-art tools: IsoQuant v. 3.3.1 (Prjibelski *et al*., 2023), StringTie v. 2.2.3 (Pertea *et al*., 2015) and TALON v. 6.0 (Wyman *et al*., 2019). For FLIC we used following parameters: “--make_downsampling --extra_filter_iso --splice_sites splice_sites_tair10_1.tsv”. Parameters for IsoQuant v. 3.3.1 were: “--complete_genedb --data_type pacbio --transcript_quantification unique_only”. FLIC and IsoQuant include mapping step a part of the pipeline; two other tools require bam files as an input so before running them we performed mapping using minimap2 v. 2.24 with parameters “-ax splice -k14 -uf -t 50 -G 10k”. After that StringTie and TALON were run with default settings.

### 2.2.17 Assessment of isoform reconstruction quality

Quality of isoform reconstruction was estimated only for isoforms that were detected in two replicates. An isoform was considered as correctly reconstructed if its 5’-end, splice junction sites and 3’-end were identical to ones annotated in TAIR10.1. Since FLIC and other tools differ in the definition of ends (FLIC defined them as regions and other tools as points) we converted FLIC regions to points and points found by other tools, to regions (see results for the details of the procedure)

### 2.2.18 Software availability

FLIC tool is available at GitHub via link: https://github.com/albidgy/FLIC and at Figshare (<u>doi: 10.6084/m9.figshare.25469317</u>). The scripts for benchmarking and comparison with the reference annotation are available at https://github.com/albidgy/FLIC_article.

## 3 Results

### 3.1 Sequencing data and preprocessing

In order to provide a framework for testing and validation of the algorithm we obtained a dedicated set of long-read RNA-seq data of leaves of *Arabidopsis thaliana* – plant model object with well-characterized transcriptome, including in terms of splice sites (Klepikova *et al*., 2016). In order to ensure representation of 3’ and 5’-ends we used a modified version of SMART protocol for cDNA synthesis and amplification and subsequent filtration of reads based on the presence of amplification primers at both ends of the read (Kasianova *et al*., 2024), for details see Supplementary Figure 1). Using PromethION P2 platform we obtained 26.8M long reads; more than 70% of reads was retained after filtration (Supplementary Table S1). As soon as an important part of FLIC algorithm is the ability to infer transcription start regions, we also generated a complementary dataset from the same samples using Cap Analysis of Gene Expression (CAGE) protocol (Murata *et al*., 2014) and compared the results.

### 3.2 Principle of FLIC algorithm

Here we present FLIC — Full-Length Isoform Constructor, a tool for reconstruction of isoforms from transcription start site (TSS) to polyadenylation site (PA). FLIC has a modular structure (Fig. 1). The main source of information on isoform structure are long cDNA reads (since FLIC was tested on ONT reads, we will further refer to them as ONT, though we expect that it is suitable for PacBio reads as well). In addition, FLIC can use several types of additional data: a set of splice junction coordinates (for example ones inferred from short-read data) and genome annotation.

**Figure 1.**
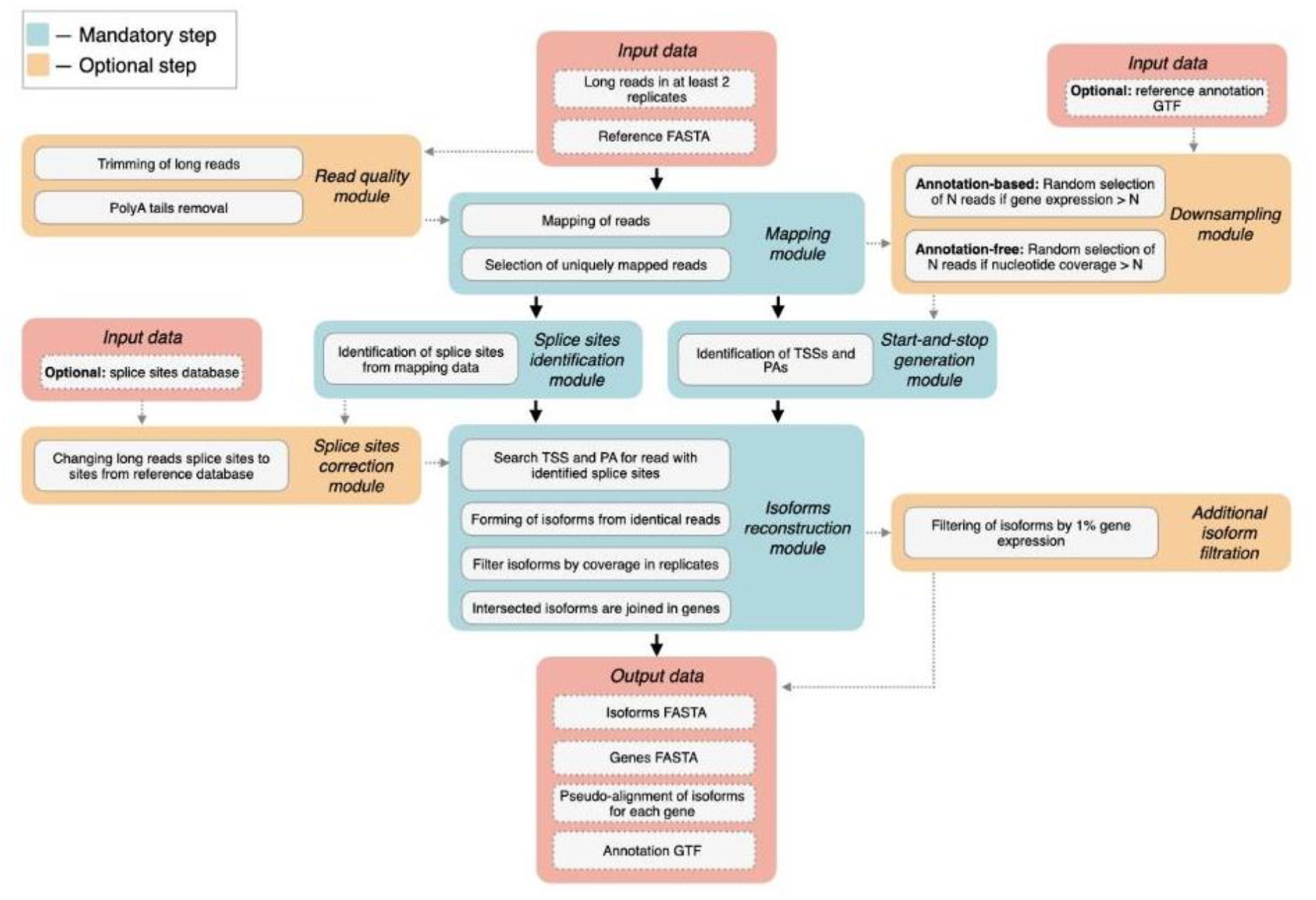
The overview of FLIC tool modules. Red blocks represent input and output files, blue blocks describe mandatory stages of the analysis, while orange blocks are optional.

First, *read quality module* takes as input ONT reads, preferably filtered as described above, and trim reads by quality score (quality threshold by default is set to 7) together with polyA tail removal.

Next, *mapping module* takes trimmed ONT reads and reference genome assembly and applies minimap2 (Li, 2018) in a splice mode to align the reads. Only uniquely mapped reads are stored for the subsequent analysis.

*Splice sites identification module* describes each read in terms of prolonged alignment gaps potentially corresponding to introns. The optional *splice sites correction module* performs splice site correction since long read mapping is prone to imperfect alignment near gaps resulting in incorrect detection of splice site coordinates (Tang *et al*., 2020; You *et al*., 2022). Thus, *splice sites correction module* uses external information on splice junction coordinates (e.g. obtained with short read mapping (Dobin *et al*., 2013) or extracted from genome annotation) to verify splice sites obtained from ONT reads. If a prolonged gap in ONT read alignment is located within +/- 10 bp range of a splice site in the predefined coordinate set, predefined splice site would be included in read description instead of initial gap coordinates. If a gap does not match any external-identified splice site, it would be considered as a deletion and discarded.

Next two modules are aimed to reconstruct TSSs and PAs of isoforms. Optional *downsampling module* (Fig. 2) deals with highly expressed genes: for genes that produce large amount of RNA molecules the absolute number of long reads derived from tattered RNA is high, this can result in incorrect TSS identification. To resolve the issue, a fixed number (1 000 by default) of reads is randomly selected from all (> 1 000) reads mapped on a gene. This step can be done using a genome annotation, if it is available, or can be run in an annotation-free mode. In the latter case at first step the coverage by uniquely mapped reads is calculated and compared with the threshold value (1000 by default). For regions for which the coverage exceeds this threshold, reads are randomly removed in an iterative way until all the positions in this region have coverage smaller than the threshold (for details see Supplementary Fig. 2).

**Figure 2.**
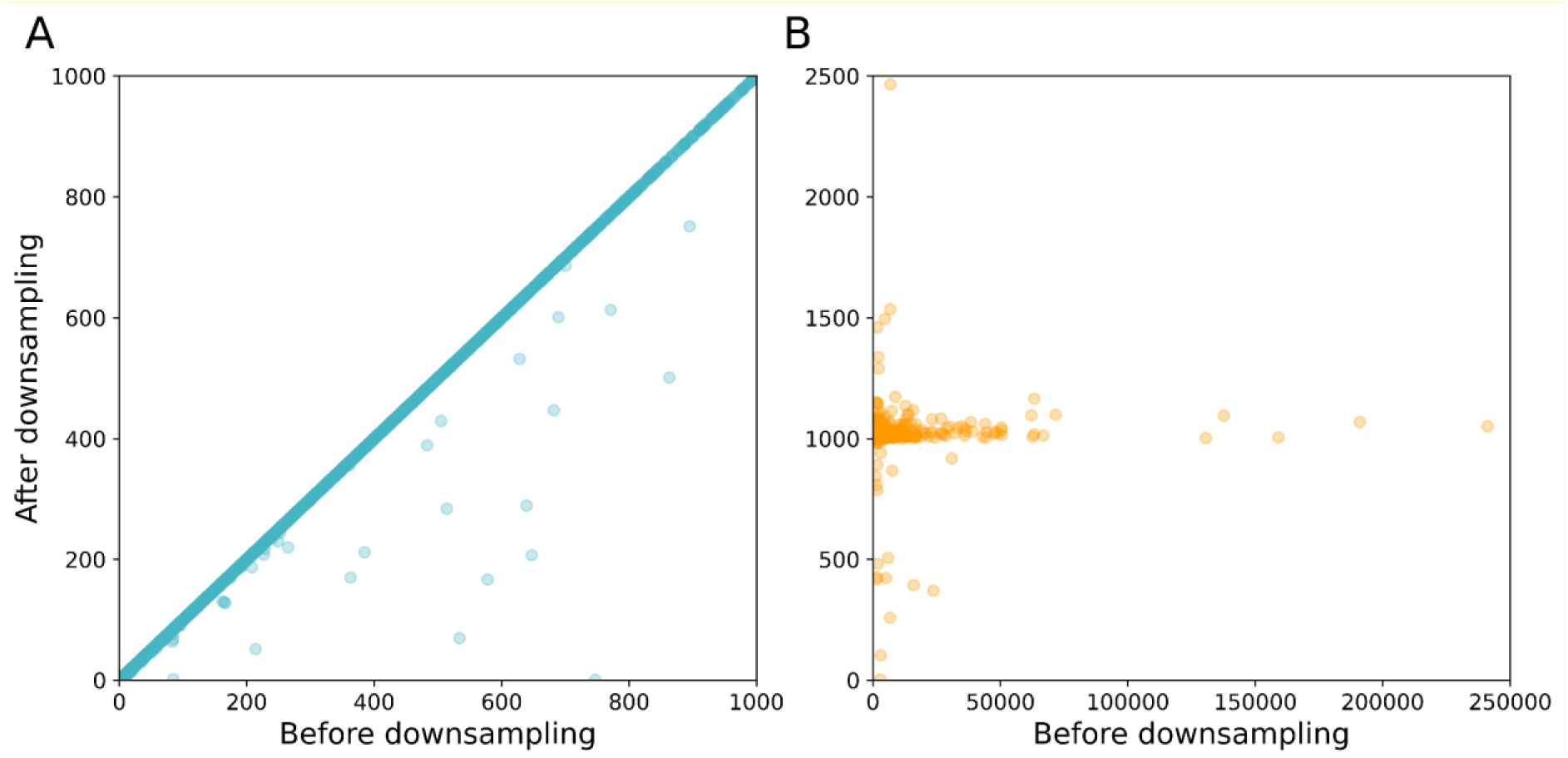
Effect of the downsampling module on gene expression in annotation-free mode. (A) Genes with ≤ 1 000 reads remain mostly unchanged after downsampling. (B) Highly expressed genes are reduced to 1 000 reads, which mainly ensures uniform coverage.

*Start-and-stop generation module* is based on peak calling. Mapped reads (either downsampled or not) are used for identification of 5’-end coordinates (presumable TSSs) and 3’-end coordinates (presumable PAs). R package CAGEfightR (Thodberg *et al*., 2019) is used for identification of regions corresponding to TSSs and PAs.

*Isoform reconstruction module* searches each read (described with splice site coordinates by *splice sites identification* and, optionally, *splice sites correction modules*) against a list of TSSs and PAs identified by *start-and-stop generation module*. Reads with identical TSS region, set of splicing sites and PA region are grouped together and form an isoform. The isoform is considered valid if it is formed by at least five reads in one biological replicate and supported by at least one read in at least one another replicate. Isoforms inferred at this step are compared by coordinates with each other. If they intersect by at least 80% they are considered as corresponding to the same gene. The borders of the gene are defined as rightmost and leftmost coordinate of all isoforms corresponding to a gene. If genome annotation is provided, reconstructed isoforms are named with existing gene IDs if they intersect with a gene by at least 80%. As output FLIC provides reconstructed isoforms and genes in FASTA format along with pseudo-alignment of isoforms for each gene and annotation in GTF format.

### 3.3 Testing and validation of FLIC on *Arabidopsis thaliana* long read transcriptome dataset

19 744 953 filtered ONT reads (5-7 million reads per replicate, for details see Supplementary Table S1) were processed by the FLIC tool. The *mapping module* of the FLIC tool returned 91-95% of reads uniquely mapped on TAIR10.1 *Arabidopsis* genome assembly (Supplementary Table S1). As FLIC can use an external set of splice junctions (e.g. inferred from exact Illumina reads) we used Illumina-based *Arabidopsis* transcriptome map (Klepikova *et al*., 2016); see Methods) as a source of information on splice junctions. The full list of splice sites detected in the transcriptome map included 254 309 unique events and 38-45% of them were identified in each replicate of leaf sample (Supplementary Table S2). ONT read alignments harbored 10.1-14.4M of prolonged gaps which potentially represent introns. We demonstrated that the vast majority of prolonged gaps in ONT reads perfectly match intron coordinates originated from Illumina data (Supplementary Table S2), though on average 454 thousand prolonged gaps per replicate were corrected or excluded by the *splice sites correction module* of FLIC (Supplementary Table S2).

At the next steps we identified TSSs and PAs. We applied *downsampling module* (to each replicate separately); 3.6-4.4% of expressed genes had coverage higher than 1 000 ONT reads (Supplementary Table S3) and were subjected to read downsampling. Next, we called peaks using *start-and-stop generation module* and obtained 117 033 TSSs and 60 246 PAs in total. Concerning the number of isoforms, FLIC results in 65 079 reconstructed isoforms belonging to 10 686 genes when applying a threshold of five or more reads supporting the isoform in at least one replicate and at least one read supporting the isoform in any other replicate. To check the reproducibility in biological replicates we reconstructed isoforms with FLIC for each replicate separately (Fig. 3A). FLIC results demonstrated high concordance of biological replicates. Next, FLIC filtered isoforms by the frequency of occurrence: for each gene only the isoforms expressed at >1% of the major isoform expression were retained resulting in 45 563 isoforms in *Arabidopsis* leaf data. We compared the subset of isoforms (ones that differ by splice sites only, not by TSS and PA positions and do not include single-exon isoforms) with the most recent *A. thaliana* genome annotation (TAIR10.1). 57% of isoforms inferred by FLIC are found in the annotation (Supplementary Fig. S3). Both sets of isoforms – generated by FLIC and annotated in TAIR10.1 – however have a large unique part – 7 165 in FLIC set and 22 605 in annotation set (Supplementary Fig. S3). This is expected because, from one side, the sample analyzed using FLIC represented only mature leaves and thus carried only leaf-specific isoforms while a plenty of unique isoforms are present in other organs, developmental stages and conditions. From the other side, TAIR10.1 annotation (which contains 59 893 isoforms identified in multiple studies on different plant organs and environmental conditions) obviously does not include all existing isoforms because many new splice junctions and isoforms were discovered in high-depth sequencing data (Klepikova *et al*., 2016; Zhang *et al*., 2022) but not included in the current annotation. Thus we did not expect the good agreement between annotation-based and FLIC-reconstructed isoforms. Isoforms reconstructed by FLIC resembled the reference isoforms in terms of intron per isoform number, isoform length, mean intron and exon number (Fig. 3B). The majority of reconstructed isoforms were shorter than 5 000 bp (only 221 isoforms exceeded the threshold).

**Figure 3.**
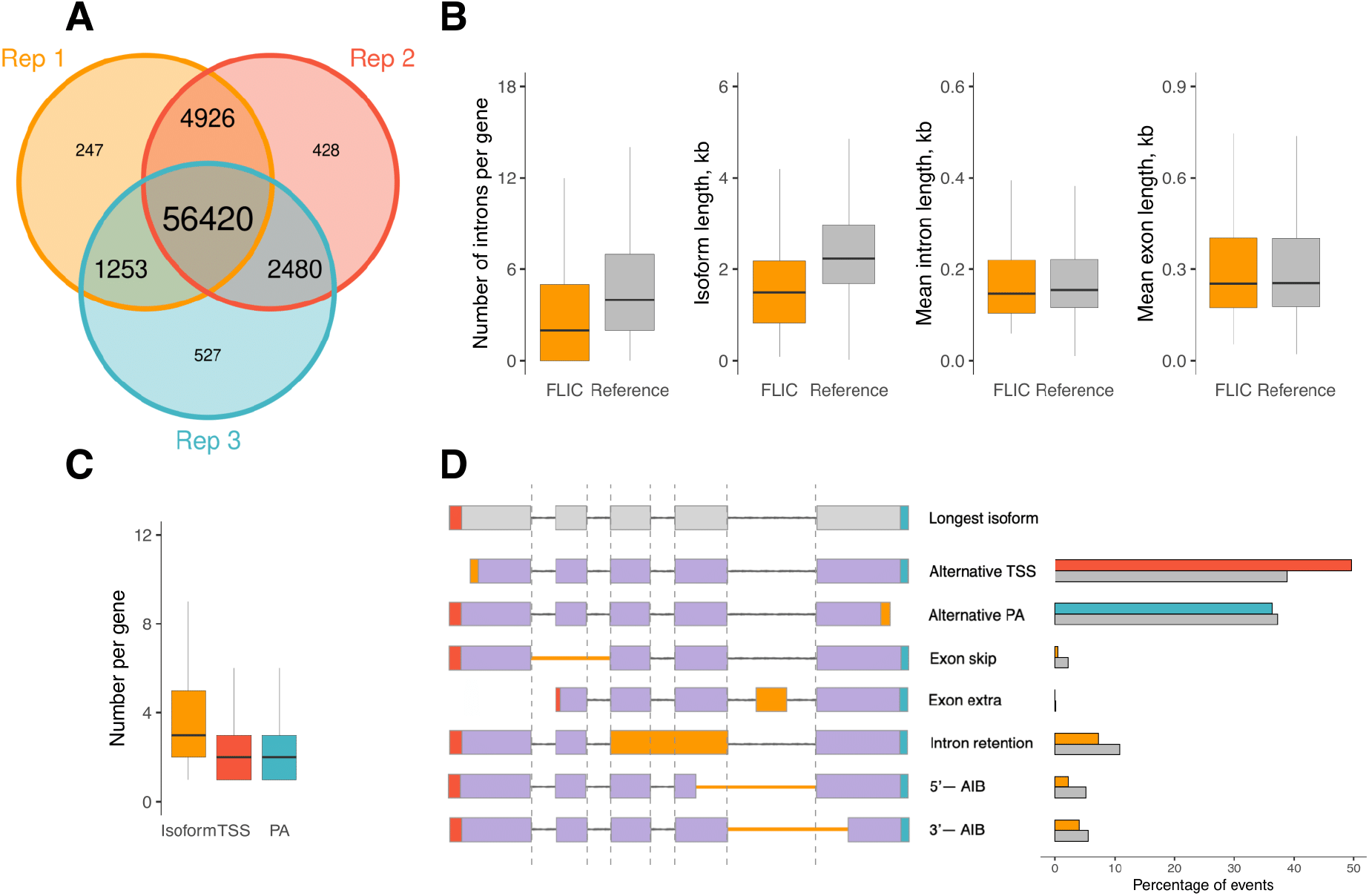
The consistency of isoforms reconstructed with FLIC. (A) The concordance of isoforms reconstructed by FLIC in each replicate of Arabidopsis leaves separately. (B) The comparison of isoform characteristics in FLIC results and genome annotation (only genes reconstructed by FLIC were selected). Outliers are not displayed. (C) The diversity of isoform number, alternative TSS and PAs in reconstructed genes. Outliers are not displayed. (D) The percentage of splicing events in reconstructed isoforms and corresponding genes in the genome annotation (grey bars). AIB - alternative intron boundary.

FLIC also has the functionality of identification of genes based on the intersection of reconstructed isoforms. The genes identified by FLIC show high congruence with ones from the annotation (Supplementary Fig. S4). The majority (75%) of genes had no more than five isoforms with median value three (Fig. 3C). 22% of genes had a single isoform in *Arabidopsis* leaf. 75% of genes had three or less TSSs and PAs (Fig. 3C), though TSSs demonstrated greater diversity: TSSs number per gene was up to 28, while maximal number of PAs – 12. About 45% of genes expressed in leaves had single TSS or PAs. Concerning the type of alternative splicing events, we found alternative TSS and PAs to be the most abundant event, followed by intron retention and the difference in 3’-boundary of intron. This is congruent with earlier reports (Martín *et al*., 2021); the distribution of alternative splicing events in the current annotation is also similar (Fig. 3D).

In contrast to most tools for isoform discovery which represent TSS and PA sites as a point, FLIC identifies them as a region through peak calling procedure. TSSs and PAs were of comparable length with median 15 and 16, respectively (Fig. 4A).

**Figure 4.**
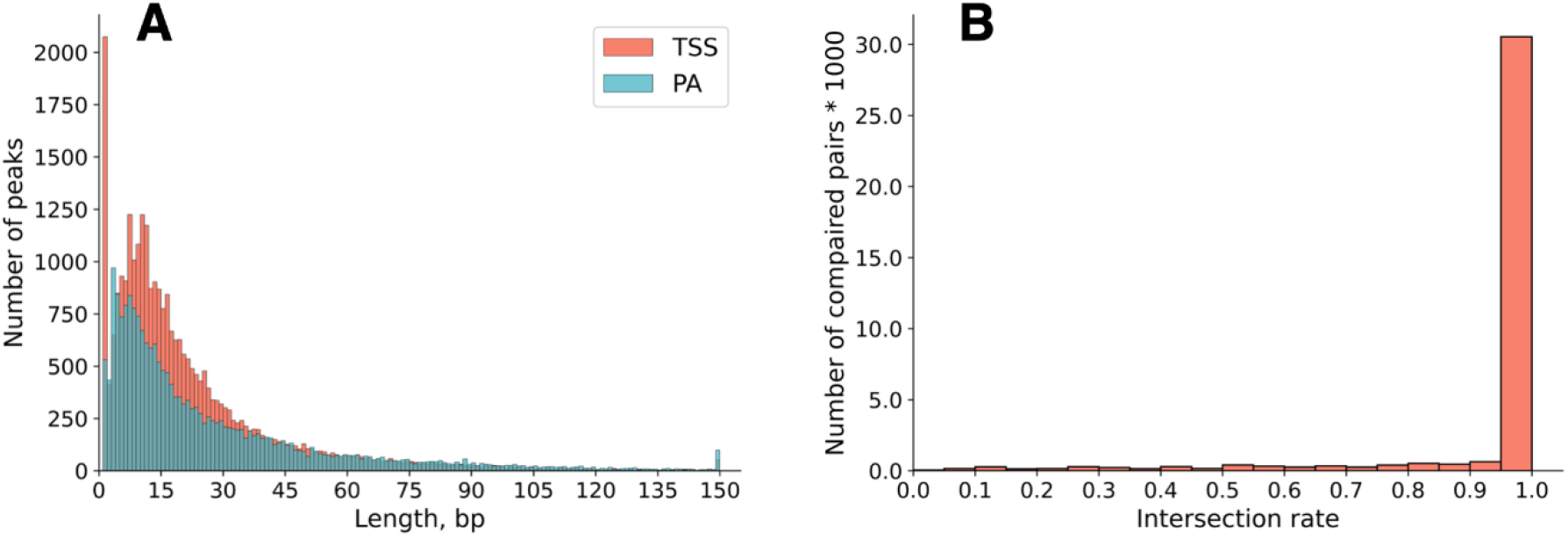
Characteristics of TSSs and PAs. (A) The distribution of TSS and PA length. TSSs and PAs longer than 150 bp were reduced to 150 bp. (B) The intersection of TSS identified by FLIC on ONT reads with TSS discovered in CAGE data. The intersection rate was determined as the length of intersected with CAGE TSS part of ONT-based TSS normalized on the length of ONT TSS.

### 3.4 Prediction of transcription start regions: comparison with CAGE

To test the validity of TSS regions identified by FLIC on ONT reads we used the results obtained by CAGE method. CAGE protocol relies on the pull-out of capped mRNAs (Shiraki *et al*., 2003) and is currently a golden standard for the identification of TSS. In order to provide an unbiased test for the performance of FLIC we performed CAGE protocol on the same RNA samples that were used for ONT sequencing (130M of high-quality reads in total, see Supplementary Table S1 for details). Using CAGE data TSS regions were identified with CAGEfightR (Thodberg *et al*., 2019) using the same parameters as in FLIC *start-and-stop generation module*. As CAGE method is prone to detection of noisy peaks outside the gene start (Georgakilas *et al*., 2020), for the purposes of this comparison we took into account only regions identified in +/-500 bp proximity of annotated starts of *Arabidopsis* genes. FLIC tool detected 46 828 TSSs using ONT reads, while 108 960 TSSs were found by CAGE. This is presumably due to much higher amount of sequencing data (40-50 M for CAGE versus 6.9-9.5M for ONT).

The number of genes with identified TSS regions was close between two data types (20 438 for CAGE and 14 967 for ONT), and 71% of genes with CAGE-based TSSs also had at least one TSS identified with ONT data. For this part of genes we found 77% of ONT-based TSSs to be intersected with TSSs found in CAGE data (Fig. 4B). 68% of not intersected ONT-based TSSs laid in a close proximity of CAGE-based TSSs (the median distance between regions was 7 bp) and only 5.3% of TSSs identified by FLIC were not supported by CAGE. The distribution of TSS regions inferred from CAGE and ONT is similar, with ONT peaks been narrower (average region size by ONT 7.1 b.p., average size by CAGE 8 b.p.).

### 3.5 Benchmarking of tools for isoform reconstruction

To compare the efficiency of isoform reconstruction, we benchmarked IsoQuant, StringTie, and TALON (Prjibelski *et al*., 2023; Pertea *et al*., 2015; Wyman *et al*., 2019). We used the *A. thaliana* annotation (TAIR10.1) as a standard. 50 059 transcripts, corresponding to 31 346 protein-coding genes (50559 transcripts and 30033 genes after additional filtering that left only those transcripts that had CAGE peaks), were considered as etalon transcripts and used to simulate PacBio reads (∼13 million reads per replicate in three replicates). Since a key advantage of FLIC is its ability to function without or with imperfect annotation, we modeled three scenarios: 1) use with no annotation provided, 2) use with perfect annotation (TAIR10.1), and 3) use with imperfect annotation. For the latter, we artificially distorted the TAIR10.1 annotation by shifting the boundaries of etalon transcripts by 100 nucleotides in various ways (Fig. 5). The programs being compared exhibit different conceptualizations of transcript boundaries. IsoQuant, StringTie, and TALON define them as single points, whereas FLIC considers them as regions corresponding to TSS and PA peaks. To ensure comparability, we conducted two types of comparisons. Firstly, we developed a procedure that allows standardizing boundary definitions as points for all programs. To achieve this for FLIC, which defines borders as peaks, not as points, we identified the peak maxima for 5’- and 3’-ends and treated them as the boundary points. Secondly, we defined boundaries as the peak regions. As all programs other than FLIC do not infer peaks for 5’- and 3’-ends, we applied the following procedure: 1) inferred peaks by FLIC using simulated reads and etalon annotation 2) identified maximal length of the peak for 5’- and 3’-ends for each gene (the largest value across all values for all isoforms of this gene), let it be N 3) for each isoform of a certain gene set the peak interval within the positions N/2 nucleotides upstream and downstream of the start and end points of this gene identified by the corresponding program. The correspondence between inferred isoforms and genes was found based on the coordinates of the isoforms (if an isoform coordinates had >10% intersection with gene coordinates, it was considered as corresponding to this gene). It should be noted that such way of comparison gives a handicap to other programs because for larger peaks the probability to overlap with the ends of etalon annotation is higher. Despite that, FLIC demonstrated the highest F1-score in two out of three comparisons (see below).

**Figure 5.**
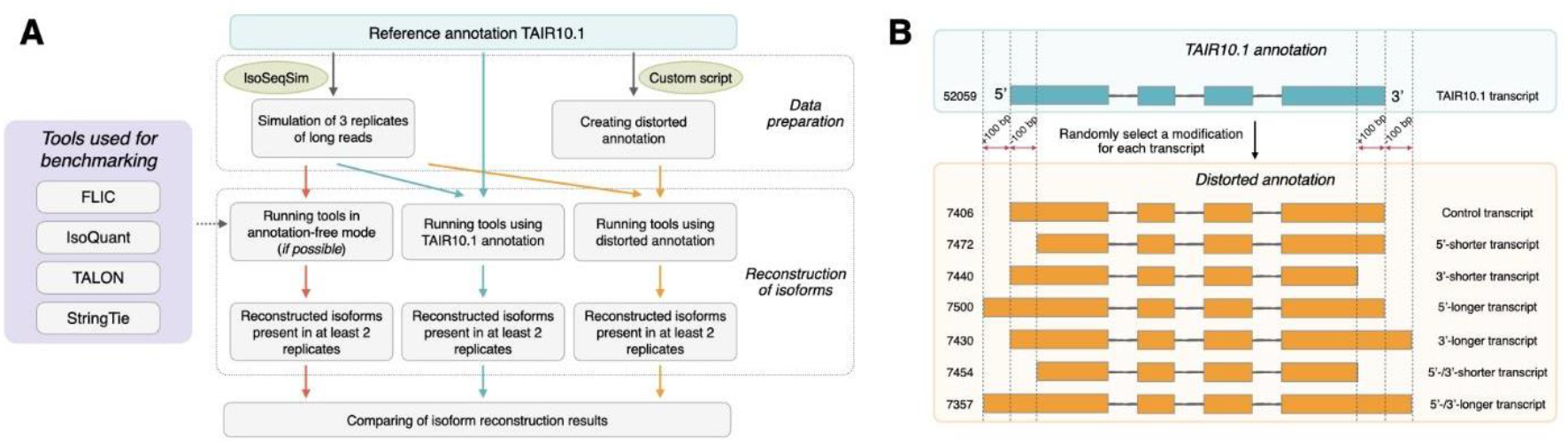
(A) Overview of benchmarking process. (B) Principles of generation of distorted annotation. Figures at the left denote the number of transcripts of each type.

When boundaries were defined as points and the unmodified TAIR10.1 annotation was used, TALON and IsoQuant exhibited the highest F1-scores (97.2% and 88.9%, respectively), while FLIC showed a comparatively lower value (66.7%). However, these values underwent significant changes when using a distorted annotation or the annotation-free mode. With the distorted annotation, the F1-score for FLIC remained stable at 66.8%, whereas for TALON and IsoQuant, it dropped dramatically to 13.8% and 14.9%, respectively (Fig. 6A). This observation demonstrates that FLIC exhibits significantly lower dependence on the annotation compared to IsoQuant and TALON. IsoQuant and TALON appear to correct their predicted boundaries based on the reference annotation, leading to a substantial performance decline under conditions of imperfect annotation. Conversely, FLIC, which utilizes the annotation only for downsampling reads from highly expressed genes, maintains stable performance.

**Figure 6.**
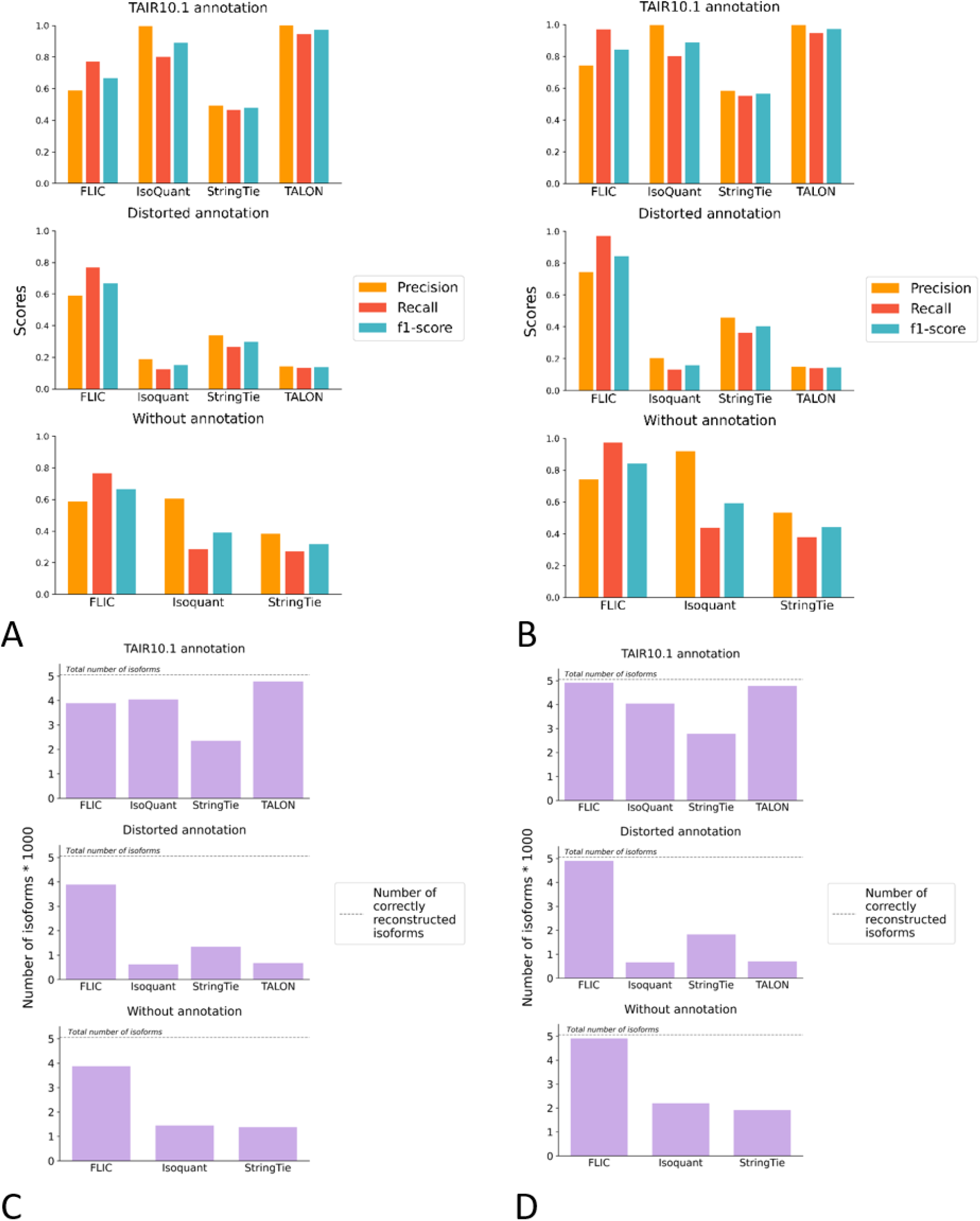
Comparison of the performance of FLIC and other tools for isoform reconstruction. (A) Precision, recall and F1-score for FLIC, IsoQuant, Talon and StringTie when 5’- and 3’-ends are considered as points. Upper panel – programs run with TAIR10.1 annotation, middle panel – with modified annotation, lower panel – in annotation-free mode. (B) Precision, recall and F1-score for FLIC, IsoQuant, Talon and StringTie when 5’- and 3’-ends are considered as peaks. Upper panel – programs run with TAIR10.1 annotation, middle panel – with modified annotation, lower panel – in annotation-free mode. (C) The number of correctly identified isoforms when 5’- and 3’-ends are considered as points. Upper panel – programs run with TAIR10.1 annotation, middle panel – with modified annotation, lower panel – in annotation-free mode. (D) The number of correctly identified isoforms when 5’- and 3’-ends are considered as peaks. Upper panel – programs run with TAIR10.1 annotation, middle panel – with modified annotation, lower panel – in annotation-free mode.

Since FLIC, IsoQuant, and StringTie possess the functionality to reconstruct isoforms without relying on an annotation, we conducted a comparison of their performance in an annotation-free mode. For FLIC, the F1-score remained stable at 66.5%, demonstrating consistent performance across all compared modes. Notably, IsoQuant exhibited a higher F1-score (38.9%) in the annotation-free mode compared to its performance with the modified annotation. Furthermore, IsoQuant demonstrated nearly identical precision to FLIC (60.6% vs. 58.7%). This observation further emphasizes IsoQuant’s reliance on the provided annotation, suggesting that the absence of annotation can lead to improved performance compared to the presence of unreliable annotation. StringTie displayed the lowest results in the annotation-free mode. When isoform boundaries were considered as TSS and PA peaks, all of the aforementioned patterns persisted, although the overall quality of isoform prediction significantly improved. For FLIC, the F1-score increased to 84%, maintaining stability across all modes (Fig. 6B).

De novo predictions of transcripts are always a trade-offs between false negatives and false positives. In terms of the number of correctly identified isoforms (true positives), FLIC outperforms the competitors in all but one comparison (Fig. 6C, D). When TAIR10.1 annotation is provided and transcript ends are defined as points, TALON identified highest number of correct transcripts, followed by IsoQuant and then FLIC (Fig. 6C). However with modified annotation and in annotation-free mode this number drops significantly, being 2-4 times lower for competitors than for FLIC. Furthermore with the distorted annotation IsoQuant and TALON predominantly reconstruct isoforms whose boundaries remained unchanged (control group), emphasizing their dependence on the accuracy of the provided annotation (Supplementary Tables S4 and S5). When ends are considered as peaks, FLIC shows highest performance, being able to correctly identify almost all transcripts (∼49 100 out of 50 559). TALON and IsoQuant follow with a small gap. The high performance of FLIC is retained in distorted annotation and annotation-free mode while for other programs it drops (Fig. 5D). Thus, FLIC consistently demonstrates its ability to provide high-quality predictions independent on an annotation provided.

## 4 Discussion

Third generation NGS platforms (Pacific Biosciences, PacBio and Oxford Nanopore technologies, ONT) have proven to be a valuable instrument for the characterization of full-length transcripts. However, many problems that hinder their application, especially for the discovery of new isoforms are still persistent. They stem from both experimental (the presence of truncated sequences, transcriptional noise, low coverage for isoforms with low expression level) and computational (mis-alignments around splice sites, high error rate) shortcomings. These problems were addressed in different ways in such tools as StringTie, FLAIR, TALON, IsoQuant (Pertea *et al*., 2015; Tang *et al*., 2020; Wyman *et al*., 2019; Prjibelski *et al*., 2023). They differ in a variety of details, in particular, the way of identification of terminal positions of isoforms, the mitigation of sequencing errors, coverage thresholds and the treatment of experimental replicates.

We present here a new approach that is centered on experimental data. Its main features are 1) the congruence of biological replicates as a criterion of quality; 2) the use of peak calling for the inference of transcription start regions and polyadenylation sites 3) the ability to work with imperfect annotation or without annotation 4) the ability to use the external information on splice sites that come from the high-depth short read data.

Short-read data (implying standard polyA-selected RNA-seq) have limited efficiency in the reconstruction of the whole isoforms but are well-suited for the detection of splice junctions (Chhangawala *et al*., 2015). Such data are available for most model objects, in form of transcriptome maps – the complex of RNA-seq experiments that cover as many as possible developmental stages, organs, tissues and conditions (e.g. (Klepikova *et al*., 2016; Stelpflug *et al*., 2016; Penin *et al*., 2021; Bullones *et al*., 2023; Sreedasyam *et al*., 2023). For non-model organisms they can be generated in a time- and cost-efficient manner. The use of external splice junctions allows to overcome the problem of generation of false isoforms stemming from inaccurate alignments of error-prone reads. This problem is less acute with HiFi PacBio reads and ONT reads generated by newer versions of sequencing chemistry that has accuracy close to 99% (https://nanoporetech.com/platform/accuracy) but is still topical for older versions.

The use of biological replicates and the estimation of their congruence is an inevitable part of gene expression analysis (see e.g. (Schurch *et al*., 2016). For splicing analysis using long reads it has not yet become a standard practice (Li *et al*., 2020; Wu *et al*., 2023; Kiyose *et al*., 2022) and even when the replicates are included in the experiment design, the presence of an isoform in replicates is not used as a criterion for the classification of it as true or false. We argue that this is an important evidence that allows to get rid of artifactual isoforms generated due to the stochastic noise of the enzymatic reactions (either in vivo, in the cell, or during sample processing and sequencing). In FLIC we require the isoform to be present in two replicates (out of three, in case of higher number of replicates this can be adjusted). This dramatically decreases the number of isoforms retained, minimizing the risk of false positives. From the other side, this leads to false negatives corresponding to low-expression level isoforms however the increase of sequencing depth helps to overcome this issue. Indeed, the results of simulation experiment show that under high coverage (> 10 million reads per replicate) it is feasible to identify almost all transcripts of the etalon annotation (FLIC correctly identifies 49 thousands out of 50,5 thousands).

The identification of terminal positions of a transcript – which, in case if full-length transcript is sequenced, correspond to transcription start and polyadenylation sites – is a challenging task. Most currently available tools for isoform discovery do not include this step, dealing with objects with well-characterized TSS and PA – such as human for which a wealth of data is available in frame of such projects as ENCODE and FANTOM. IsoQuant defines terminal positions as the outermost positions of the read mapped to the reference. This is a valid approach however it is susceptive to artifacts such as transcription noise, in particular, the read-through of RNA polymerase (which is not uncommon, (Caldas *et al*., 2024). In FLIC the recognition of transcription start and polyadenylation regions is done using the approach of peak calling, similar to the one used for methods specifically aiming at characterization of 5’ and 3’-ends (CAGE, PAS-seq).

Another important feature of FLIC, that was clearly highlighted by the benchmarking done in different modes with regard to the annotation, is its independence from the reference annotation. While competitor tools showed superior performance when the etalon annotation is provided, they had much less success in annotation-free mode and when modified annotation (the model for imperfect annotation) is used. While the development of sequencing technologies and corresponding bioinformatics tools has provided a toolkit for efficient assembly, even for large and complex eukaryotic genomes (Cosma *et al*., 2022; Mochizuki *et al*., 2023; Kong *et al*., 2023), genome annotation is still lagging. The imperfection of genome annotation is virtually universal for any group of eukaryotes (see e.g. (Park *et al*., 2023; Vuruputoor *et al*., 2023). Long full-length transcriptome reads can be a game changer in this problem as they offer a great opportunity for the evidence-based annotation on transcripts, including 5’- and 3’-UTRs. However, current state-of-the-art tools such as BRAKER3 (Gabriel *et al*., 2024) do not specifically incorporate long reads focusing instead on the assembly of short reads using StringTie. This gap is addressed in our pipeline. Taken together we expect that FLIC be a useful tool for isoform analysis in a wide range of species, not only ones that have well-characterized transcriptome, such as human. Also, it has the potential to provide de novo annotations for non-model species for which genomes are just being sequenced.

## Data availability

The ONT and CAGE sequencing reads generated in this study have been submitted to the NCBI BioProject database (https://www.ncbi.nlm.nih.gov/bioproject/) under accession number PRJNA1087576.

## Supplementary figures

**Figure S1** The workflow of ONT library construction and ONT reads filtration.

**Figure S2** Downsampling module workflow in annotation-free mode.

**Figure S3**The intersection of isoforms obtained by FLIC and presented in Arabidopsis genome annotation for genes expressed in leaf sample

**Figure S4** The comparison of gene length reconstructed by FLIC and presented in Arabidopsis genome annotation

## Supplementary tables

**Table S1** Mapping statistics of reads used in the study

**Table S2** Statistics of splice sites detection in ONT data by splice sites identification module of FLIC

**Table S3** Statistics of gene number downsampled by downsampling module of FLIC

**Table S4** True positive (TP) rates for isoform reconstruction in the presence of distorted annotation and in annotation-free modes (pointes)

**Table S5** True positive (TP) rates for isoform reconstruction in the presence of distorted annotation and in annotation-free modes (ranges)

## Acknowledgments

Nanopore sequencing was performed using the core facilities of the Lopukhin FRCC PCM “Genomics, proteomics, metabolomics” (http://rcpcm.org/nauchnye-issledovanija/centr-kollektivnogo-polzovaniia/). We are grateful to Artem Kasianov for his work at the inception of this project, Alexey Kovtun for the name of the tool, Vsevolod Makeev and Ivan Kulakovskiy for helpful discussion. We are grateful to Sofia Prokopchuk for the algorithm tests.

## Funding

This work was supported by funding from Ministry of Science and Higher Education, project [075-15-2021-1064 - data, FFRW-2024-0003 - analysis].

## Conflict of interest

None of the authors have conflicts of interest to declare.

## Notes

https://github.com/albidgy/FLIC

### Competing Interest Statement

The authors have declared no competing interest.

