## Supplementary_Files for "Full-length isoform constructor (FLIC) – a tool for isoform discovery based on long reads"

### Supplementary Files Content

| | |
|---|---|
| **Figure S1** | The workflow of ONT library construction and ONT reads filtration. |
| **Figure S2** | *Downsampling module* workflow in annotation-free mode. |
| **Figure S3** | The intersection of isoforms obtained by FLIC and presented in *Arabidopsis* genome annotation for genes expressed in leaf sample |
| **Figure S4** | The comparison of gene length reconstructed by FLIC and presented in *Arabidopsis* genome annotation |
| **Table S1** | Mapping statistics of reads used in the study |
| **Table S2** | Statistics of splice sites detection in ONT data by splice sites identification module of FLIC |
| **Table S3** | Statistics of gene number downsampled by *downsampling module* of FLIC |
| **Table S4** | True positive (TP) rates for isoform reconstruction in the presence of distorted annotation and in annotation-free modes (pointes) |
| **Table S5** | True positive (TP) rates for isoform reconstruction in the presence of distorted annotation and in annotation-free modes (ranges) |

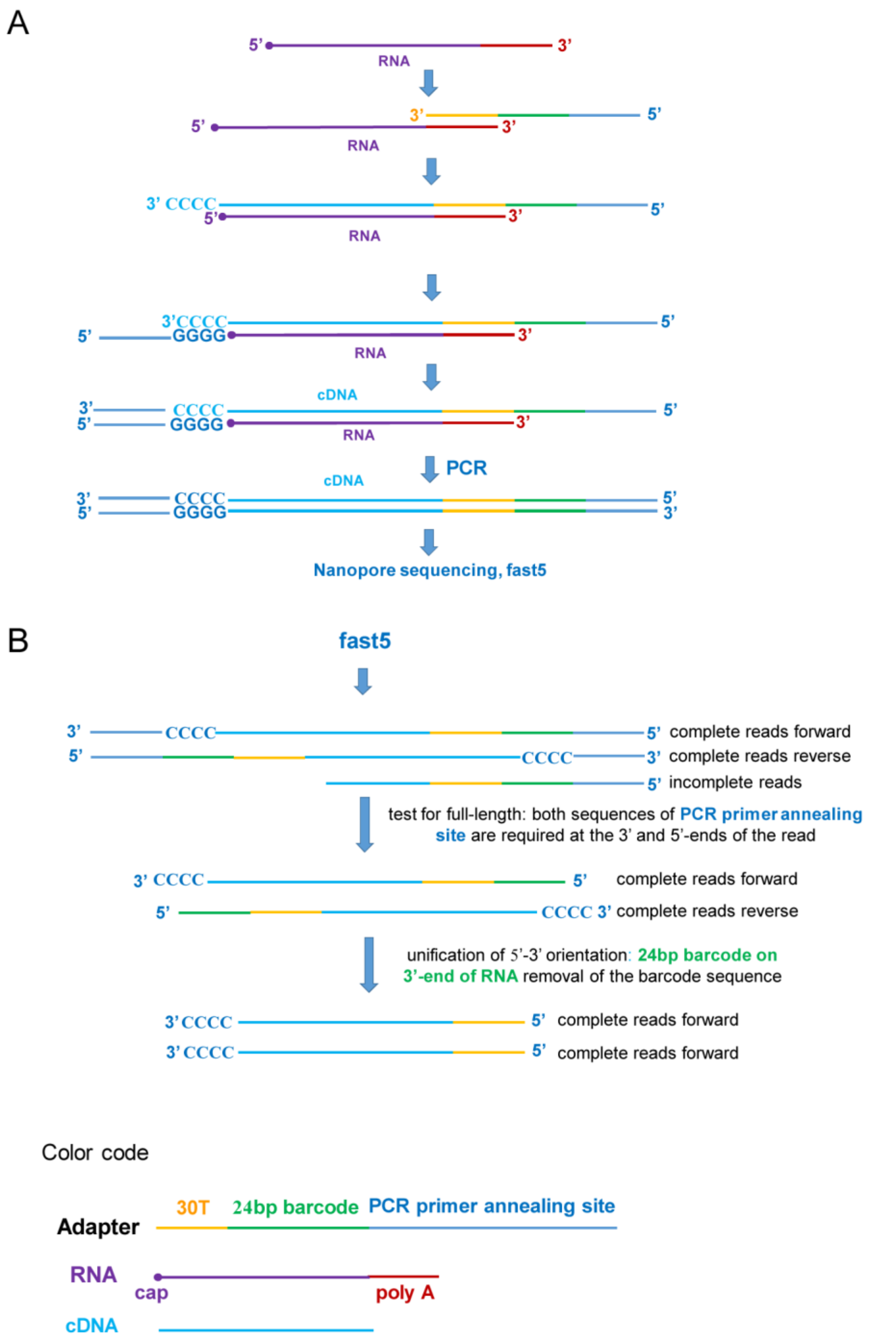


**Supplementary Figure 1.** The workflow of ONT library construction and ONT reads filtration. **(A)** The adaptation of SMART protocol. **(B)** Stages of ONT reads filtration.

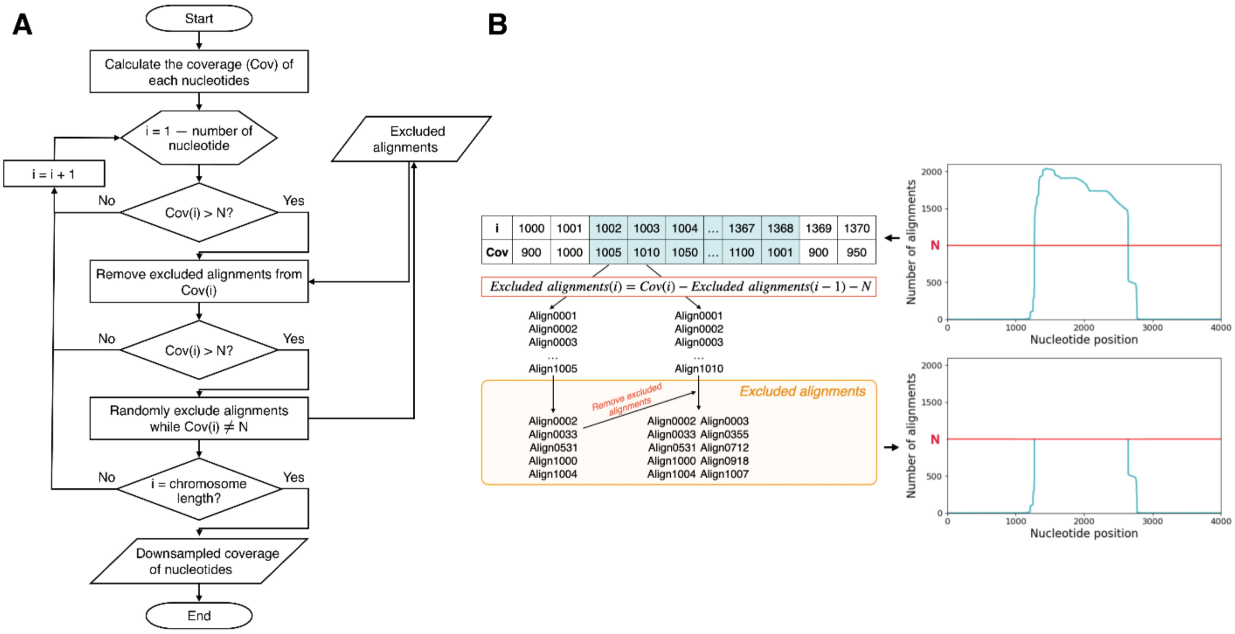


**Supplementary Figure 2.** *Downsampling module* workflow in annotation-free mode. **(A)** Flowchart of the module's operation. **(B)** Conceptual illustration showing the module's functionality during the downsampling process.

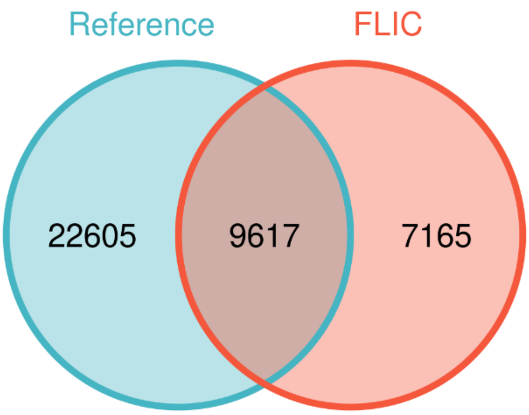


**Supplementary Figure S3**. The intersection of isoforms obtained by FLIC and presented in *Arabidopsis* genome annotation for genes expressed in leaf sample. TSSs and PAs were excluded from the analysis.

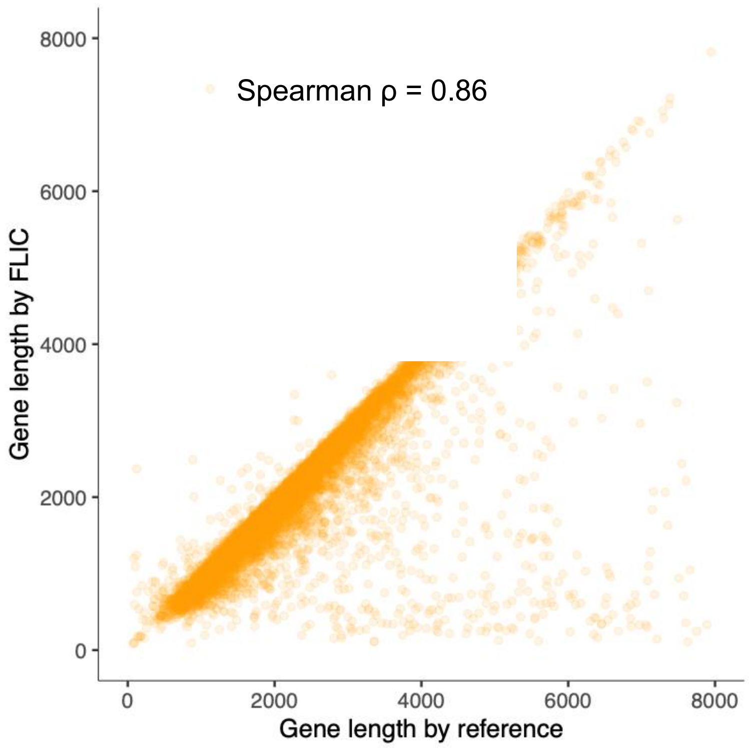


**Supplementary Figure S4**. The comparison of gene length reconstructed by FLIC and presented in *Arabidopsis* genome annotation.

**Supplementary Table S1. Mapping statistics of reads used in the study**

**Three replicates of *Arabidopsis* leaves, ONT libraries**

| | Replicate 1 | Replicate 2 | Replicate 3 |
|---|---|---|---|
| Raw read number | 9 459 558 | 10 389 060 | 6 952 504 |
| Number of read trimmed by qualitry and M1 primers presence (percentage of raw read number) | 7 057 521 (75%) | 7 405 646 (71%) | 5 281 786 (76%) |
| Number of mapped reads (percentage of trimmed reads) | 6 969 873 (99%) | 7 284 457 (98%) | 5 200 431 (98%) |
| Uniquely mapped reads (percentage of trimmed reads) | 6 681 390 (95%) | 6 742 474 (91%) | 4 985 049 (94%) |

**Three replicates of *Arabidopsis* leaves, CAGE libraries**

| | Replicate 1 | Replicate 2 | Replicate 3 |
|---|---|---|---|
| Raw read number | 50 177 822 | 45 187 079 | 40 547 826 |
| Number of read trimmed by qualitry (percentage of raw read number) | 48 491 378 (97%) | 42 463 180 (94%) | 38 909 790 (96%) |
| Number of mapped reads, primary alignments only (percentage of trimmed reads) | 47 215 840 (97%) | 41 363 934 (97%) | 37 910 884 (97%) |

**Supplementary Table S2. Statistics of splice sites detection in ONT data by splice sites identification module of FLIC**

| | Replicate 1 | Replicate 2 | Replicate 3 |
|---|---|---|---|
| Number of reference splice sites identified in ONT reads (percentage of total number (254 309) of splice sites in Illumina data | 112 623 (44.3%) | 114 608 (45.1%) | 97 477 (38.3%) |
| Total number of prolonged gaps in ONT reads | 14 915 516 | 14 769 888 | 10 423 633 |
| Number of ONT gaps perfectly matched with reference splice sites (percentage of total number) | 13 803 535 (92.5%) | 13 645 115 (92.4%) | 9 730 327 (93.3%) |
| Number of ONT gaps intersected with reference splice sites (percentage of total number) | 877 525 (5.9%) | 876 276 (5.9% | 596 516 (5.7% |
| Number of ONT gaps not matched with reference splice sites (percentage of total number) | 234 456 (1.6%) | 248 497 (1.7%) | 96 790 (0.9%) |

**Supplementary Table S3. Statistics of gene number downsampled by downsampling module of FLIC**

| | Replicate 1 | Replicate 2 | Replicate 3 |
|---|---|---|---|
| Total number of genes covered by at least five ONT reads | 16 810 | 17 322 | 16 250 |
| Number of genes for which downsampling was applied (percentage of total number) | 719 (4.3%) | 761 (4.4%) | 577 (3.6%) |
| Number of genes without downsampling (percentage of total number) | 16 091 (95.7%) | 16 561 (95.6%) | 15673 (93.4%) |

**Supplementary Table S4. True positive (TP) rates for isoform reconstruction in the presence of distorted annotation and in annotation-free modes. TSSs and PAs were treated as points (nucleotide position with maximal coverage inside corresponding range)**

| Type of gene annotation disturbance | Number of genes | FLIC, % TP | Isoquant, % TP | StringTie, % TP | TALON, % TP | FLIC anno-free, % TP | Isoquant anno-free, % TP | StringTie anno-free, % TP |
|---|---|---|---|---|---|---|---|---|
| Unchanged genes | 7 177 | 77.2 % | 79.4% | 46.7% | 94.3% | 76.9 % | 28.4% | 26.9% |
| 5' shortage | 7 264 | 76.3 % | 3.1% | 10.6% | 0.0% | 76.5 % | 29.1% | 27.4% |
| 3' shortage | 7 232 | 76.8 % | 1.0% | 20.3% | 0.0% | 77.2 % | 29.3% | 27.9% |
| 5' elongation | 7 274 | 76.3 % | 0.0% | 35.7% | 0.0% | 75.9 % | 28.4% | 27.3% |
| 3' elongation | 7 217 | 77.4 % | 0.0% | 27.1% | 0.0% | 76.9 % | 28.5% | 27.3% |
| 5' and 3' shortage | 7 233 | 76.3 % | 3.4% | 20.2% | 0.0% | 76.1 % | 28.8% | 26.8% |
| 5' and 3' elongation | 7 162 | 77.9 % | 0.0% | 25.7% | 0.0% | 77.6 % | 28.3% | 27.7% |

**Supplementary Table S5. True positive (TP) rates for isoform reconstruction in the presence of distorted annotation and in annotation-free modes. TSSs and PAs were treated as ranges.**

| Type of gene annotation disturbance | Number of genes | FLIC, % TP | Isoquant, % TP | StringTie, % TP | TALON, % TP | FLIC anno-free, % TP | Isoquant anno-free, % TP | StringTie anno-free, % TP |
|---|---|---|---|---|---|---|---|---|
| Unchanged genes | 7 177 | 97.2% | 79.4% | 55.1% | 94.2% | 97.2% | 43.7% | 37.2% |
| 5' shortage | 7 264 | 96.4% | 6.3% | 13.6% | 1.8% | 96.8% | 44.3% | 37.4% |
| 3' shortage | 7 232 | 96.8% | 1.3% | 27.2% | 0.1% | 97.3% | 43.6% | 38.4% |
| 5' elongation | 7 274 | 97.2% | 0.7% | 47.7% | 1.7% | 97.2% | 43.2% | 38.0% |
| 3' elongation | 7 217 | 97.1% | 0.0% | 41.5% | 0.1% | 97.1% | 43.0% | 37.6% |
| 5' and 3' shortage | 7 233 | 97.1% | 4.9% | 28.2% | 0.1% | 97.2% | 44.1% | 37.7% |
| 5' and 3' elongation | 7 162 | 97.4% | 0.0% | 39.7% | 0.1% | 97.4% | 43.1% | 38.2% |
